# Spatiotemporally Mapping Thermodynamics of Lysosomes and Mitochondria using Cascade Organelle-Targeting Upconversion Nanoparticles

**DOI:** 10.1101/2022.02.16.480800

**Authors:** Xiangjun Di, Qian Peter Su, Dejiang Wang, Yongtao Liu, Mahnaz Maddahfar, Jiajia Zhou, Dayong Jin

**Affiliations:** Institute for Biomedical Materials & Devices (IBMD), Faculty of Science, University of Technology Sydney, Sydney, NSW 2007, Australia; School of Biomedical Engineering, Faculty of Engineering and Information Technology, University of Technology Sydney, Sydney, NSW 2007, Australia; UTS-SUStech Joint Research Centre for Biomedical Materials & Devices, Department of Biomedical Engineering, Southern University of Science and Technology, Shenzhen, China

**Keywords:** Lysosome, Mitochondria, Nanothermometry, Upconversion Nanoparticles (UCNPs)

## Abstract

The intracellular metabolism of organelles, like lysosomes and mitochondria, are highly coordinated spatiotemporally and functionally. The activities of lysosomal enzymes significantly rely on the cytoplasmic temperature, and heat is constantly released by mitochondria as the byproduct of ATP generation during active metabolism. Here, we develop temperature-sensitive LysoDots and MitoDots to monitor the *in situ* thermodynamics of lysosomes and mitochondria. The design is based on upconversion nanoparticles (UCNPs) with high-density surface modifications to achieve the exceptionally high sensitivity of 2.7% K^-1^ and accuracy of 0.8 K for nanothermometry to be used in living cells. We show the measurement is independent of the intracellular ion concentrations- and pH values. With Ca^2+^ ion shock, the temperatures of both lysosomes and mitochondria increased by 2∼4 °C. Intriguingly, with Chloroquine treatment, the lysosomal temperature was observed to decrease by up to ∼3 °C, while mitochondria remained relatively stable. Lastly, with oxidative phosphorylation inhibitor treatment, we observed a 3∼7 °C thermal increase and transition from mitochondria to lysosomes. These observations indicate different metabolic pathways and thermal transitions between lysosomes and mitochondria inside HeLa cells. The nanothermometry probes provide a powerful tool for multi-modality functional imaging of subcellular organelles and interactions with high spatial, temporal and thermal dynamics resolutions.

**Graphical Abstract:** Cascade organelle-targeted nano-thermometers based on upconversion LysoDots and MitoDots.

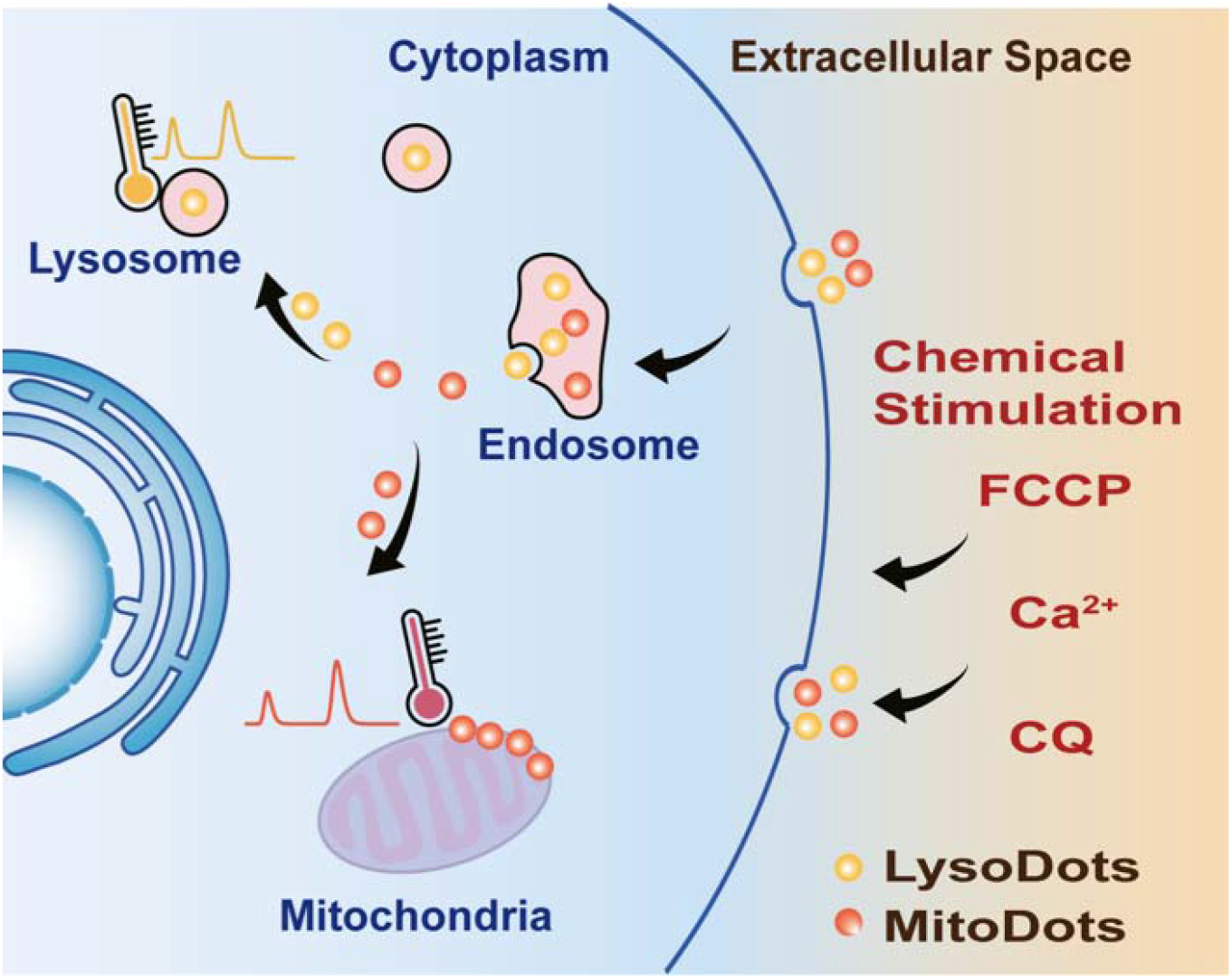

## Introduction

The intracellular temperature and its dynamics are vital to maintaining the homeostasis of organelles as well as their biochemical reactions and metabolic processes. Thermal variations indicate whether cells are under their healthy physiological or disease status [1, 2]. The coordination of different organelles is required to maintain intracellular homeostasis and normal cellular functions. The intracellular organelles not only possess their specific functions but also communicate with each other through membrane contacts or membrane fusion, which together contribute to the survival, growth and division of the cell. Lysosomes and mitochondria, as the two major contributors to enzyme activity and energy production, are essential to cellular metabolism participating in many key biological processes such as autophagy, proliferation, cell death, etc. [3-5]. Dysfunction of lysosomes and mitochondria has been found in several diseases [6].

Both lysosomes and mitochondria’s maintenance of their physiological functions and cellular homeostasis reply on intracellular thermodynamics [7-9]. Lysosomes are highly dynamic organelles responsible for the turnover of some proteins and lipids through digestive enzymes [10, 11]. The enzymatic activities are highly dependent on a healthy pH and temperature environment. During the process of heatstroke, the pH values of lysosomes were observed to increase, which leads to cell death. A higher or lower temperature may reduce the enzymatic activities and further disturb the cellular homeostasis [12]. Mitochondria are involved in the cellular respiratory and function as the energy factory [13]. During respiratory activity, mitochondria transform energy from carbohydrates to ATP, simultaneously releasing heat as a byproduct.

Due to the lack of enabling techniques and tools, the functions of lysosomes and mitochondria have been independently studied in the past, though there must be intense crosstalks between the two organelles [3]: The dysfunctional mitochondria lead to the elevation of lysosomal pH [14]; and in some lysosomal storage diseases, the defects in the lysosomes also contribute to the dysfunction of mitochondria because the abnormal mitochondria cannot be cleared by lysosomes, leading to the pathological signalling [15]. At present, the interactions of lysosomes and mitochondria are usually studied from either their direct physical membrane contacts using super-resolution microscopy [16] or signalling pathways using molecular biology tools [3]. It is unclear about the spatiotemporal thermodynamics of lysosomes and mitochondria during the pathological process to understand how they maintain homeostasis within a live cell.

Up to now, a series of organic and inorganic nanoscale thermometers have been reported [2, 17]. Organic fluorescent dyes can efficiently target organelles but suffer from the limited sensitivities and the issue of photobleaching. Inorganic nano-thermometers are bright with good photostability and chemical stability but large in size and require careful modifications to improve their delivery efficiency in living cells and targeting specificity to the organelles, which challenges their accurate detection of the local temperature of a specific organelle. Lanthanide-doped upconversion nanoparticles (UCNPs) show great potential as nano-thermometers, due to their high brightness, superior photochemical stability and excellent temperature-responsive optical properties with large anti-Stokes shifts and long luminescence lifetimes [18]. Though many advances have been made in using UCNPs-based nano-thermometers to monitor the temperature variations in living cells, tissues, or animals [19, 20], organelle-specific temperature sensing remains challenging in the physiological environment.

Here, we report that surface modification of high-density polymer linkers can assist UCNPs to be specifically guided and accumulated into lysosomes and mitochondria in live cells. We establish a novel cascade targeting strategy in this work. We first introduce PEGMEMA_80_-*b*-EGMP_3_ di-block copolymers to transfer the hydrophilic UCNPs to hydrophilic nanoparticles that can be internalized through the endocytosis process and ended in the lysosomes [21]. We name the structure of UCNPs@copolymer as **LysoDots**. To facilitate the escape of UCNPs from lysosomes and relocation to pass through the potential barrier of mitochondrial membrane, we then introduce Poly-L-Lysine (PLL) to further functionalize UCNPs with a mitochondrial targeting moiety with high lipophilicity, (3-carboxypropyl)triphenylphosphonium bromide (TPP) UCNPs [22]. We name the structure of UCNP@PLL@TPP as **MitoDots**. This strategy allows two sets of nano-thermometers with organelle-targeted properties to simultaneously and quantitatively sense the *in situ* thermodynamics of lysosomes and mitochondria. The series of LysoDots and MitoDots demonstrates a sensitivity of 2.7% K^-1^ and an accuracy of 0.8 K in HeLa cells, and more importantly, their performances are independent of the ion concentrations or pH conditions. We have observed intriguing spatiotemporal thermodynamics in HeLa cells: i) with organelle non-specific Ca^2+^ shock, both lysosomal and mitochondrial temperature increase 2-4 °C; ii) with lysosome-specific drug treatment, the lysosomal temperature drops ∼3 °C while mitochondria remain 37 °C; iii) with mitochondria-specific drug treatment, we observed an interesting temperature increment (3-7 °C) and thermal transition (1-2 min delay) from mitochondria to lysosomes. These observations indicate different metabolic pathways and thermal transitions between lysosomes and mitochondria. The live-cell thermal probes provide a powerful toolbox with multimodality and multi-functional imaging capacities, including spatial, temporal and thermal dynamics of selective organelles with high resolution and accuracy.

### Design, Synthesis and Characterization of Organelle-Targeted Nano-Thermometers

Following our previously established protocols [23-25] to construct a stable and biocompatible UCNPs based nanothermometer, PEGMEMA_80_-*b*-EGMP_3_ di-block copolymers were used to transfer hydrophobic UCNPs into hydrophilic ones, and UCNPs@copolymer showed long-term colloidal stability in the culture medium (**Figure S1A**). In this design, the phosphate acid anchoring groups offer greater affinity to lanthanide ions exposed on the surface of UCNPs than oleic acid, and the carboxylic acid group at the terminal of the di-block copolymer allows further modification and conjugation. In this work, to improve the bio-conjugation efficiency to a mitochondria-targeted molecule, (3-carboxypropyl)triphenylphosphonium bromide (TPP) [21], Poly-L-Lysine (PLL) was introduced to enrich the density of amine groups on the surface of UCNPs@copolymer through attractive electrostatic interactions. By the carbodiimide reaction, UCNPs@PLL were further crosslinked with TPP to form UCNPs@PLL@TPP that exhibited chemical stability in the culture medium (**Figure S1B**). The three kinds of modified UCNPs (UCNPs@copolymer, UCNPs@PLL and UCNPs@PLL @TPP) showed good morphology uniformity and monodispersity. The size of UCNPs increased from 29.97 ± 0.98 nm of the as-synthesized UCNPs to 30.23 ±1.48, 30.54 ± 1.17, and 31.57 ± 1.23 nm (measured by TEM in **Figure 1B**). The dynamic light scattering (DLS) results (**Figure 1C**) confirmed the high uniformity with the average values of hydrodynamic size increasing from 55.67 ± 0.62 nm to 59.80 ± 0.25 nm and 66.51 ± 1.49 nm after each step of surface modifications. The Zeta potential results (**Figure 1D**) indicated the successful modifications of each step, as the surface charge turned from negative 14.28 ± 0.59 mV to positive 20.77 ± 0.30 mV with the exposure of PLL and 12.18 ± 0.47 mV with TPP. The presence of PEGMEMA_80_-*b*-EGMP_3_ di-block copolymers, PLL and TPP on the surface of UCNPs were further confirmed by ATR-FTIR. As shown in **Figure S2A**, after grafting the PEGMEMA_80_-*b*-EGMP_3_ di-block copolymers on the surface of the nanoparticles, the asymmetric COO-stretching at 1547 cm^-1^ of oleic acid molecules completely disappeared, with both the characteristic ATR-FTIR spectral peak at 1100 cm^−1^ (C−O−C stretching) and the new Ph-P band at 1440 cm^-1^ confirming the success in PEGMEMA_80_-*b*-EGMP_3_ di-block copolymers and TPP ligands modifications.

**Figure 1.**
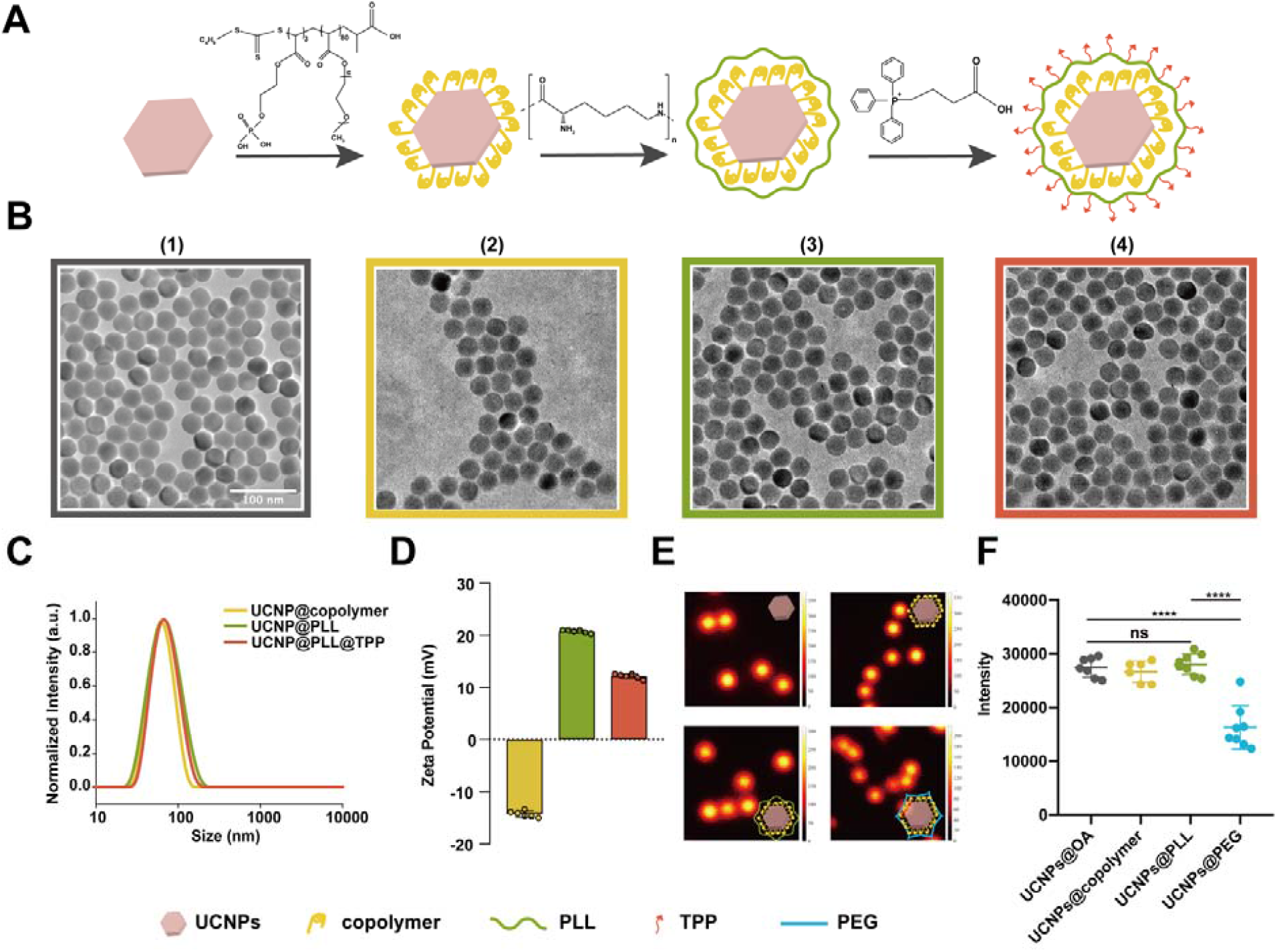
Surface modification strategy and characterization of UCNPs to produce efficient organelle-targeting nano-thermometers. **(A)** Schematic illustration of the cascade modification steps based on UCNPs with the surface modified from oleic acid (OA, pink), to PEGMEMA_80_-b-EGMP_3_ di-block copolymers (yellow), poly-l-lysine (PLL, green) and (3-carboxypropyl)triphenylphosphonium bromide (TPP, red), respectively. **(B)** Representative transmission electron microscopy (TEM) images of 1) UCNPs-OA (grey), 2) UCNPs@copolymer (LysoDots, yellow), 3) UCNPs@PLL (green), and 4) UCNPs@PLL@TPP (MitoDots, red), respectively. Scale bar, 100 nm. **(C)** Dynamic light scattering (DLS) size analysis of UCNPs after different surface modifications (colour-codes same as in Fig1B). **(D)** ζ-Potentials of UCNPs after different surface modifications (colour-codes same as in Fig1B). **(E)** Luminescence emission images from single UCNPs coated with OA, copolymer, PLL, and 4Arm-PEG-NH_2_ (PEG) under 980 nm excitation. **(F)** Statistics of the emission intensity of single individual UCNPs in Fig.1E.

### The Roles of Poly-L-Lysine (PLL)

To demonstrate the advantage of the cascade modification strategy used in this work, samples from our previous work [26], where the crosslinked polymer networks composed of PEGMEMA_80_-*b*-EGMP_3_ di-block copolymers and 4Arm-PEG-NH_2_ on the surface of UCNPs (UCNPs@PEG), was compared. but relatively low emission intensity. Because the chemical bonds (O-H, C-H, or N-H) on the UCNPs surface match the photon states of the host material and influence the non-radiative relaxation of the excited lanthanide ions [24]. As shown in **Figure 1E & F** and **Figure S2B**, the intensity measurements indicated that the fluorescent intensity of UCNPs@PLL was 1.8 times higher than that of UCNPs@PEG. This may be due to the chemical bonds (O-H, C-H, or N-H) on the UCNPs surface that influence the non-radiative relaxation of the excited lanthanide ions [24]. Moreover, compared with 4Arm-PEG-NH_2,_ PLL with amino acid chain provides a lot of more amine groups on the UCNPs surface. 10 mg of TPP and 5 mg of nanoparticles were conjugated overnight through EDC-NHS reaction. After the reaction, unreacted TPP was completely removed by centrifugal ultrafiltration and the supernatant was retained so that, according to the UV−vis absorptions at 268 nm of various concentrations of TPP in the supernatant (see Methods section and **Figure S2C**), the conjugation efficiency of TPP has been enhanced from 0.332 mg to 1.545 mg for 5 mg of UCNPs@PEG and UCNPs@PLL, respectively.

In our cascade organelle-targeting strategy, after being internalized by the cells [21], UCNPs@copolymer nanoparticles translocate from the early endosomes to the late endosomes that ferry the nanoparticles into lysosomes, and therefore as a nano-thermometer to naturally target lysosome (**LysoDots**); UCNPs@TPP nanoparticles first being accumulated in the lysosomes, and due to the lipophilic and positive TPP moieties, can escape from endo-lysosomes to specifically target mitochondria (**MitoDots**).

### *in vitro* Thermo-responsive Properties of LysoDots and MitoDots

The luminescence spectrum change of Yb^3+^ and Er^3+^ co-doped UCNPs strongly depends on the temperature (**Figure 2A[1]**) [27]. Within a single nanoparticle, Yb^3+^ ions transfer the sensitized 980 nm photon energy to Er^3+^ ions that upconvert the energy of multiple photons to emit two distinct temperature-responsive green emissions at 515-535 nm (centred at 525 nm) and 535-570 nm (centred at 545 nm) bands (**Figure 2A [2]**). The intensity ratio of 525 nm and 545 nm bands following Boltzmann distribution, regardless of the concentrations of nanoparticles [26],

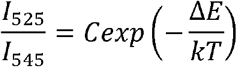

where *I*_*525*_ and *I*_*545*_ are the integrated fluorescent intensities at the 525 nm and 545 nm emission bands, respectively; *C* is a constant; Δ*E* is the energy gap; *k* is the Boltzmann constant and *T* is the absolute temperature. The calibration curve of nanoparticles in the water was plotted showing a great linear fitting (R^2^ = 0.99375, **Figure 2A[3]**).

**Figure 2.**
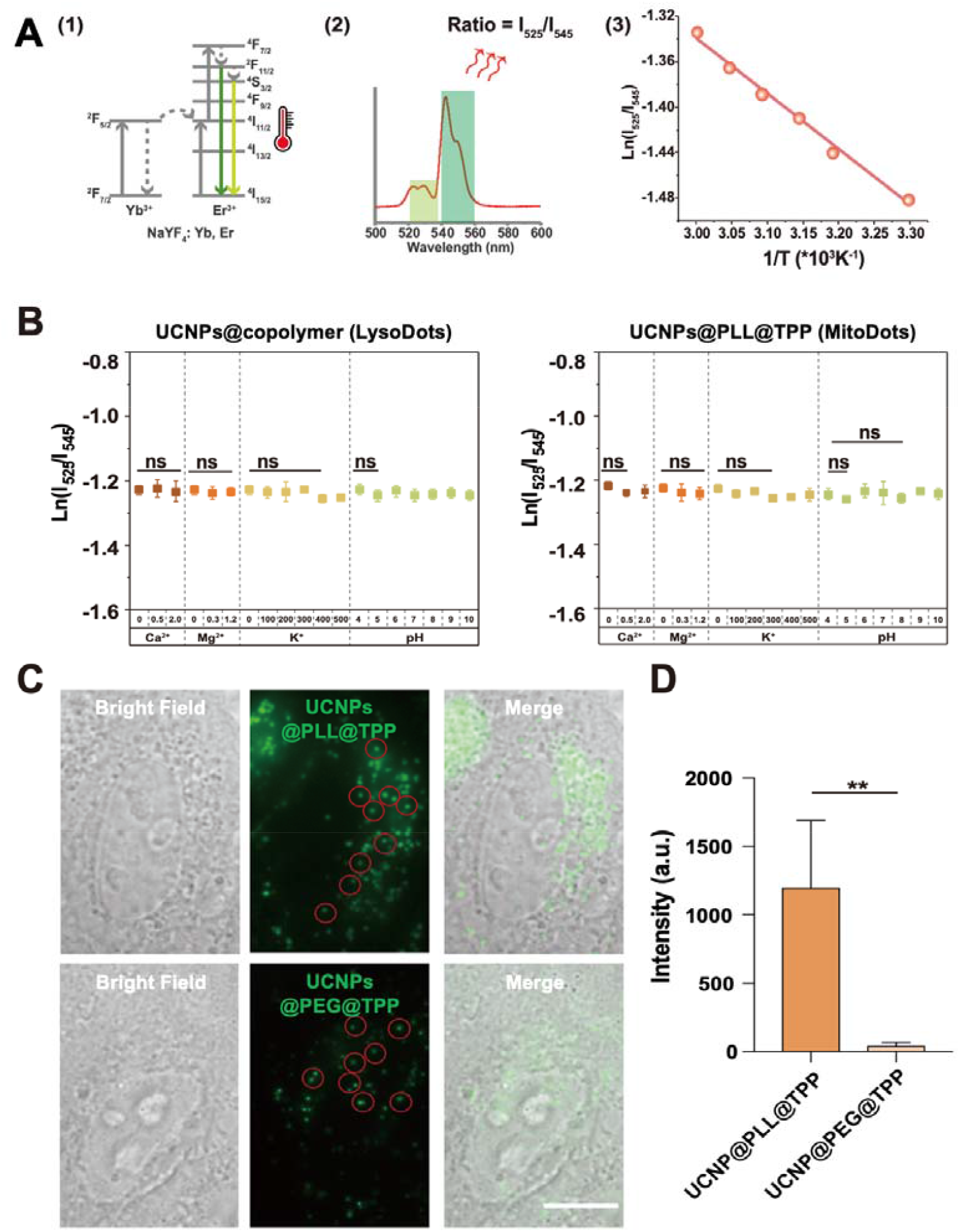
Stable and efficient LysoDots and MitoDots. **(A)** Energy level (1) and ratiometric thermometry (2) of temperature-sensitive UCNPs. Standard temperature sensing curve (3) by plotting ln(I525/I545) vs 1/T to calibrate the thermometric scale in hydrophilic solution. The equation is y = 0.12665x-0.4888 (R^2^ = 0.99375, Pearson’s R = - 0.99687). **(B)** Stability tests of ratiometric thermometry at different Ca^2+^ ion concentrations (0-2 mM), Mg^2+^ ion concentrations (0∼1.2 mM), K^+^ ion concentrations (0∼500 mM) and pH values (4∼10). N = 3 independent repeats. **(C)** Bright Field, fluorescence, and merge images of UCNPs@PLL@TPP (MitoDots) and UCNPs@PEG@TPP in HeLa cells. Scale bar = 10 μm. **(D)** Quantification of enhanced emission intensities from UCNPs@PEG@TPP to UCNPs@PLL@TPP (MitoDots) used in HeLa cells. N = 10 region of interests (ROIs).

Living cells are complex systems with the environment changing in both spatial and temporal domains. For example, the pH value is much lower in lysosomes than that in other organelles. The concentrations of Ca^2+^ in mitochondria and lysosomes also vary across the different phases of the cell cycle. To validate the thermal detection stability of LysoDots and MitoDots, we measured the temperature-dependent spectrum in different pH (4-10) and ion concentration conditions, including Ca^2+^ (0-2 mM), Mg^2+^ (0-1.2 mM) and K^+^ (0-500 mM). **Figure 2B** confirms that the luminescence responding to the temperature of LysoDots (**Figure 2B** left panel) and MitoDots (**Figure 2B** right panel) are independent of either ion concentration or pH conditions.

### Enhanced Targeting Efficiency of MitoDots

To demonstrate the advance of the cascade modification strategy in enhancing the mitochondrial targeting efficiency, both MitoDots, based on UCNPs@PLL@TPP (**Figure 2C** top panel) and UCNPs@PEG@TPP use in our previous work [26] (**Figure 2C** bottom panel) were incubated with HeLa cells for 12 hrs. As shown in **Figure 2D**, the average intensity of individual foci from UCNPs@PLL@TPP (MitoDots) was almost 10-times higher than that of UCNPs@PEG@TPP. The results suggest that the more TPP on the surface of UCNPs@PLL enabled the more efficient mitochondria-targeting ability of nano-thermometry MitoDots.

### Specificities of the Cascade Organelle-targeting and Accumulation of LysoDots and MitoDots

Using LysoTracker and MitoTracker as controls, we further performed both the independent and the simultaneous colocalization assays in HeLa cells to evaluate the intracellular specific localizations and organelle accumulation efficiencies of LysoDots and MitoDots. It is based on a purpose-built total internal reflected fluorescence (TIRF) microscopy system with 561 nm, 647 nm and 980 nm excitation lasers and a live cell incubator (see Methods section for details).

We first checked the targeting specificity of LysoDots by incubating HeLa cells with LysoDots alone for 12 hrs, followed by LysoTracker Red and MitoTracker DeepRed staining for 30 min before TIRF imaging. As shown in **Figure 3A top panel**, LysoDots (cyan) colocalized cohesively with LysoTracker (magenta) with similar dotted morphology. The Pearson’s coefficient between LysoDots and LysoTracker (**Figure 3B**) was as high as 0.95, almost twice higher than that between LysoDots and Mito-tracker (0.54). Usually, we consider a Pearson’s coefficient higher than 0.6 as credible colocalizations [28].

**Figure 3.**
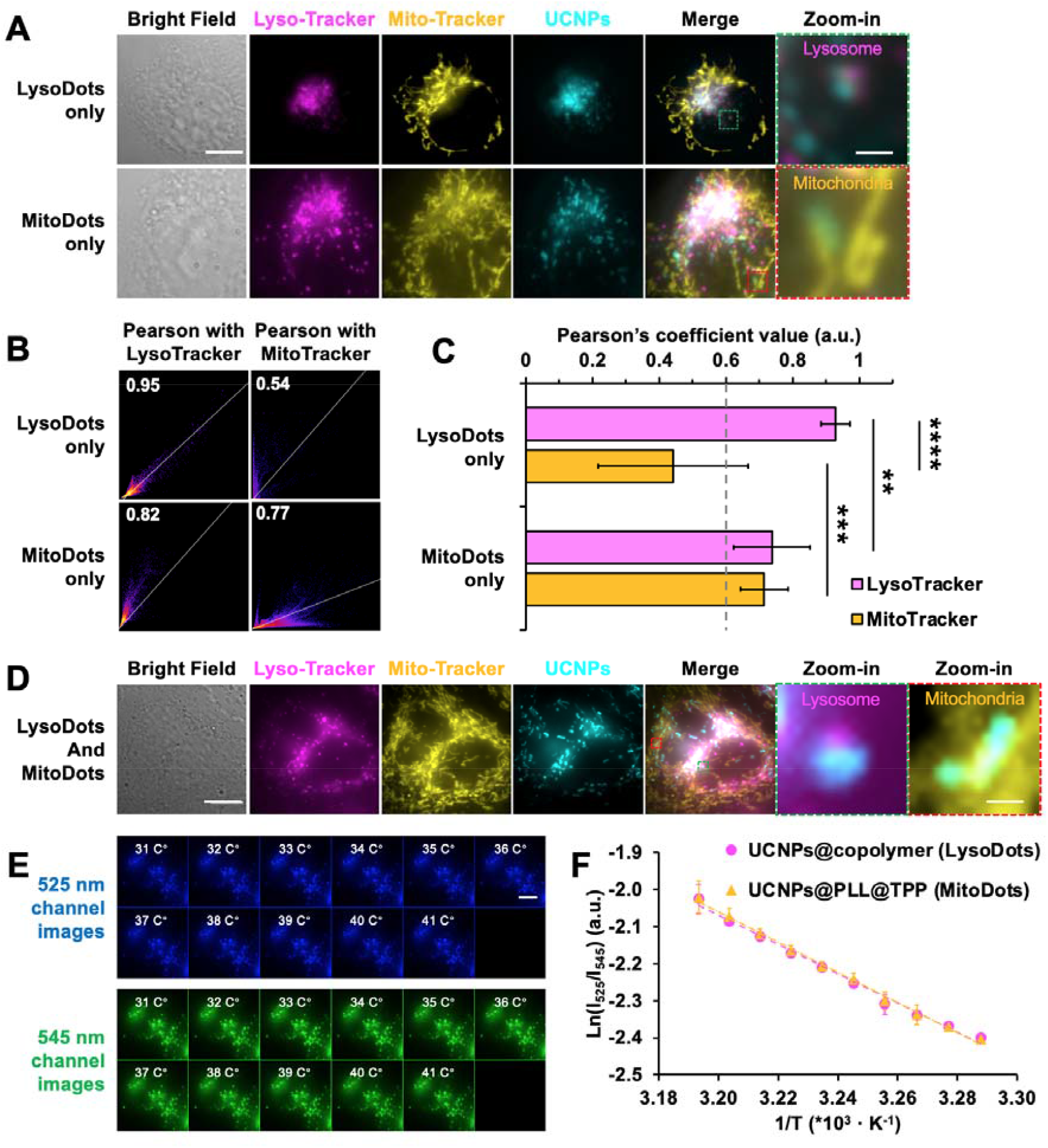
Organelle-specific accumulations of LysoDots and MitoDots. **(A)** Images of representative cells which was incubated with LysoDots (cyan, top panel) or MitoDots only (cyan, bottom panel) for 12 hrs followed by LysoTracker (magenta) and MitoTracker (yellow) staining for 30 min before fixation and TIRF imaging. The zoom-in areas show the colocalization and morphology. Scale bar = 10 μm for the whole-cell images and 1 μm for the zoom-in images. **(B)** Pearson’s coefficient values of LysoDots with LysoTracker and MitoTracker (top panel) and MitoDots with LysoTracker and MitoTracker (bottom panel). **(C)** Statistics of Pearson’s coefficient in Fig.3B. Data points represent mean ± SD. N = 10 ROIs. Significance label **, p<0.01; ***, p<0.001. **(D)** Images of a representative cell incubated with both LysoDots and MitoDots (cyan) for 12 hrs and stained with LysoTracker (magenta) and MitoTracker (yellow) 30 min before fixation and TIRF imaging, and the zoom-in areas to show the colocalization and morphology. **(E)** Snapshot images of 525 nm emission (blue, top panel) and 545 nm emission (green, bottom panel) from UCNPs when external temperature increased from 31 to 41 C°. **(F)** Plot of ln(I525/I545) v.s. 1/T to calibrate the thermometric scale of LysoDots (magenta line) and MitoDots (orange line) in HeLa cells. Data represent mean ± SD (n = 5 cells). Scale bar = 10 μm.

Similarly, we then checked the specificity of MitoDots by incubating HeLa cells with MitoDots (cyan) alone for 12 hrs, followed by LysoTracker and MitoTracker staining. As shown in **Figure 3A bottom panel**, we found that MitoDots also colocalized cohesively with MitoTracker (yellow) with similar linear and network morphologies. The Pearson’s coefficients (**Figure 3B**) between MitoDots and LysoTracker and between MitoDots and MitoTracker were 0.82 and 0.77, respectively, revealing the process of MitoDots’ escape from lysosomes to mitochondria. The significantly higher Pearson’s coefficient between MitoDots and MitoTracker (0.77), compared with that between LysoDots and MitoTracker (0.54, p < 0.001, **Figure 3C**), confirmed that the sufficient amount of TPP moieties on the surface of MitoDots has efficiently facilitated nanothermometers to anchor onto mitochondria.

To get an *in situ* calibration curve for temperature sensing, we plotted the temperature value versus the I_525_/I_545_ ratios of LysoDots and MitoDots in paraformaldehyde (PFA) fixed HeLa cells by changing the temperature using a temperature-controllable incubator. As shown in **Figure 3D**, both LysoDots and MitoDots were incubated with HeLa cells for 12 hrs, then cells were stained with LysoTracker and MitoTracker for 30 min and followed by PFA fixation before TIRF imaging when the temperature increased from 31 °C to 40 °C, both the logarithmic values of the I_525_/I_545_ ratio for LysoDots and MitoDots showed gradual and linear fluorescence relative to the reciprocal temperature (**Figure 3E**), i.e. y = 9.97417x – 3.76589 (R^2^ = 0.99685, Pearson’s r = -0.99842) and y = 10.3715x – 3.88731 (R^2^ = 0.99741, Pearson’s r = -0.9987), respectively. As shown in **Figure 3F**, the root-mean-square deviation (RMSD) of these two linear fittings was measured as 0.007025, indicating no difference between LysoDots and MitoDots. The relative sensing sensitivity at 32 ºC was 2.7% K^-1^ and the temperature resolution was ∼0.8 K.

### Spatiotemporally Mapping the Thermodynamics of Lysosomes and Mitochondria Under Chemical Stimulations

By incubating HeLa cells with both LysoDots and MitoDots for 12 hrs and with LysoTracker and MitoTracker for 30 min, we simultaneously mapped the thermodynamics of both lysosomes and mitochondria under external chemical stimulations, including both the organelle non-specific and organelle-specific treatments.

We first treated HeLa cells with Ca^2+^ ion shock, which is an organelle non-specific treatment. The Ca^2+^ ion shock can promote the pumping of ions and accelerate respiration reactions [29]. Ionomycin calcium salt is an ionophore that makes the cell membrane highly permeable for Ca^2+^ ions [30], which induces intracellular stress, possibly causing damage to both lysosomes and mitochondria in HeLa cells [29]. As shown in **Figure 4A-C**, upon a 1 μM ionomycin calcium salt treatment, the lysosomal and mitochondrial temperatures first sharply increased by 2∼3°C within 6 minutes before dropping back together in the next 20 minutes (**Figure 4B, magenta and orange lines**). Compared with the control using the solvent treatment, the temperature (**Figure 4C, grey lines**) remained at a relatively stable level regardless of the addition of 1:1000 DMSO into the culture media.

**Figure 4.**
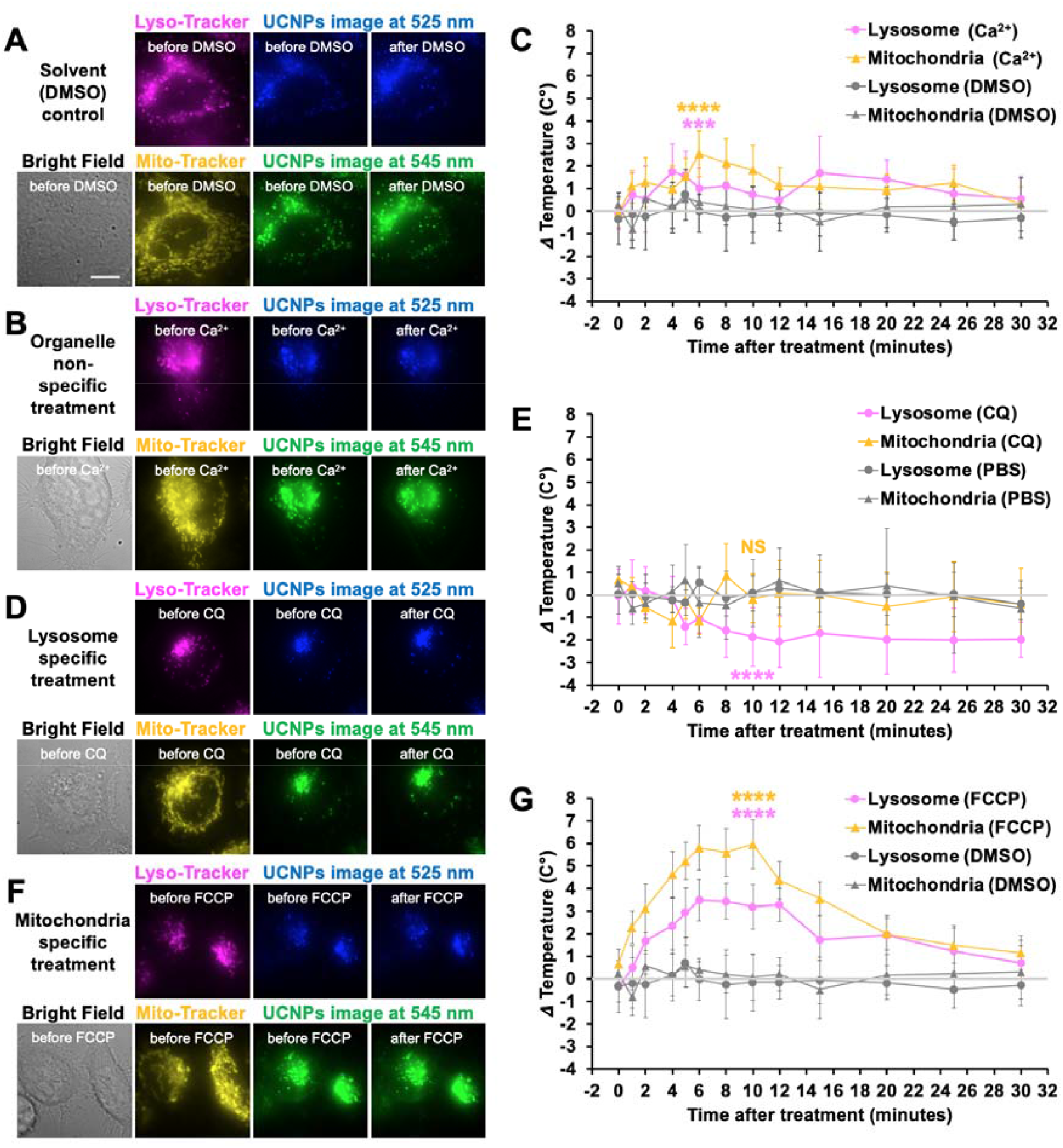
Distinct lysosomal and mitochondrial thermal dynamics in response to chemical stimulations in living HeLa cells. **(A-B)** Live-cell TIRF images of representative cells (grey for bright field) which was incubated with LysoDots and MitoDots (blue for 525 nm emission and green for 545 nm emission) for 12 hrs, stained with LysoTracker (magenta) and MitoTracker (yellow) 30 min before the imaging and treated with 1:1000 DMSO as the solvent control (**A**) and Ca^2+^ ion shock (**B**). Scale bar = 10 μm. **(C)** The thermodynamic curves of lysosome with Ca^2+^ ion shock (magenta circles), mitochondria with Ca^2+^ ion shock (orange triangles), lysosomes with DMSO (control, grey circles) and mitochondria with DMSO (control, grey triangles) within 30 min after treatment and Student’s t-test at 6 min (***, p < 0.001; ****, p < 0.0001). **(D)** Bright-field (grey), LysoTracker (magenta), MitoTracker (yellow), 525 nm (blue) and 545 nm (green) emission of UCNPs images of a representative cell before and after the treatment with 200 nM Chloroquine (CQ). **(E)** The thermodynamic curves of lysosome with CQ (magenta circles), mitochondria with CQ (orange triangles), lysosomes with PBS (control, grey circles) and mitochondria with PBS (control, grey triangles) within 30 min after treatment and Student’s t-test at 10 min (NS, no significance; ****, p < 0.0001). **(F)** Bright-field (grey), LysoTracker (magenta), MitoTracker (yellow), 525 nm (blue) and 545 nm (green) emission of UCNPs images of a representative cell before and after the treatment with 10 μM FCCP. **(E)** The thermodynamic curves of lysosome with FCCP (magenta circles), mitochondria with FCCP (orange triangles), lysosomes with DMSO (control, grey circles) and mitochondria with DMSO (control, grey triangles) within 30 min after treatment and Student’s t-test at 10 min (****, p < 0.0001).

Next, we treated live cells with a lysosome-specific Chloroquine (CQ) stimulation (**Figure 4D**). CQ has originally been used to treat malaria and is now a sensitizing agent to treat certain cancers [31, 32]. CQ is a lysosomotropic weak base, which increases the pH of lysosomes and therefore inhibits autophagic degradation in the lysosomes [33]. Intriguingly, the nanothermometry measurements by LysoDots and MitoDots revealed a significant decrease in lysosomal temperature by ∼3 °C after 10 min treatment of 200 nM CQ, while the mitochondrial temperature remains relatively stable as the basal level in HeLa cells (**Figure 4E, magenta and orange lines**). In comparison, the temperature in the solvent treatment groups (**Figure 4E, grey lines**) remained at a relatively stable level after adding PBS into the culture media.

We then treated HeLa cells with a mitochondria-specific carbonyl cyanide-4-(trifluoromethoxy)phenylhydrazone (FCCP, **Figure 4F**). FCCP is an inhibitor of mitochondrial oxidative phosphorylation, as it disrupts the ATP synthesis by transporting protons across the mitochondrial inner membrane [34]. LysoDots and MitoDots not only revealed the significant increase of both the lysosomal and mitochondrial temperature by almost 3∼7 °C in the first 10 min after adding 10 μM FCCP (P < 0.0001 by Students’ t-test, **Figure 4G, magenta and orange lines**), but also mitochondria release a large amount of heat, which is consistent with the previous observation [26], but with higher accuracy using MitoDots. A clear delay by approximately 1∼2 min in the time domain during the process of both temperature increase (0-6 min) and decrease (10-14 min) between lysosomes and mitochondria. The significant temperature increase reflects a large amount of heat released by mitochondria, with the trend being consistent with the previous observation [26]. The new ability of MitoDots suggests higher accuracy, and the observation of time delays further indicate a thermal transition from mitochondria to the lysosomes. In the following 20 min, the lysosomal and mitochondrial temperature eventually recovered to the original temperature, but the cellular morphology significantly changed, indicating cell death after mitochondrial dysfunction.

## Conclusion and Discussion

The intracellular thermodynamics, as one of the most pivotal biophysical parameters, plays a critical role in maintaining the cells’ homeostasis or driving the cells into dysfunction status [35], regulated by the coordinations of organelles and other crucial cellular activities. Our knowledge about the functions of inter-organelle communication and combinations are limited, as current tools and technologies only allow the functions of organelles to be studied as individual entities. In mammalian cells, the recycle factory Lysosomes and the energy factory of mitochondria closely interact with each other [3]: lysosomes digest macromolecules into free amino acids, sugar, and lipids for biosynthesis and energy production [10], and mitochondria make use of small molecules to produce the energy [13, 36]. From the metabolic view, mitochondria, and lysosomes communicate with each other as they coordinate the metabolites degradation, transportation, and production, which are always accompanied by ATP production and heat release. From the signalling pathway, the dysfunction of lysosomes or mitochondria will affect the function of each other leading to multiple human diseases [9, 15]. Therefore, it is desirable to visualize the thermodynamics of lysosomes and mitochondria with sufficiently high spatial and temporal resolutions.

With the advances made in non-contact luminescent nano-thermometers, LysoDots and MitoDots designed in a cascade organelle-targeting fashion can simultaneously monitor the temperature variations of lysosomes and mitochondria. Following the typical endocytosis internalizations, both types of nano-thermometers are first accumulated into endosomes and translocated to lysosomes. The large amount of TPP on the surface of MitoDots would facilitate themselves to escape from lysosomes and penetrate the potential barrier of the mitochondrial intermembrane. Therefore, the *in situ* lysosomal and mitochondrial thermodynamics under different physiological and chemical stimuli can be plotted by the highly stable and sensitive ratiometric nano-thermosensors. LysoDots and MitoDots enable us to monitor the Ca^2+^-, CQ- and FCCP-induced thermodynamics in the lysosomes and mitochondria within living HeLa cells. Such a cascade organelle-targeting platform becomes a powerful tool to visualize the interactomes between lysosomes and mitochondria as the nanoscale temperature dynamics is the key to maintaining the cellular homeostasis in living cells.

In addition to temperature, the pH value in lysosomes also affects their enzymatic activities. It has been reported that an increase in temperature causes the lysosomal pH value to be elevated [12]. Therefore, it is desirable to simultaneously monitor the pH dynamics in lysosomes during the temperature measurement. As UCNPs have been used to sense the *in situ* pH values [37], it would be feasible to develop the “all-in-one” nanoscale sensors based on LysoDots.

The ability in mapping the distinct thermodynamics highlights the extensive range of applications using the organelle-targeting strategy to study the vital biological processes, among lysosome, mitochondria, endoplasmic reticulum [38], Golgi apparatus [39], lipid droplet [16], peroxisome [16], and so on. The colocalization analysis in **Figure 3** suggests the amount of MitoDots being stuck in the lysosome after 12 hrs incubation were not negligible, additional rational design strategies, e.g. introducing another small molecule, are needed to further improve the escape efficiency of MitoDots, besides enriching the density of TPP demonstrated in this work. At least, this work suggests the cascade modification and bioconjugation strategy can target other organelles and establish a library of organelle-targetted nano-thermometers.

In this work, we observed for the first time the temperature decrease in lysosomes under CQ treatment, and the thermal transition from mitochondria to lysosomes under FCCP stimulation. To better reveal the thermal transmission across multiple organelles, it becomes critical to further increase the spatial and temporal resolutions of our imaging system [40], so that the new series of nano-thermometers can be used to map the long-term thermodynamics of individual organelles across cell cycles or even in deep tissues [41-43] with single-particle sensitivity and resolution. Introducing the state-of-the-art *in situ* organelle imaging and assay approaches, including live-cell super-resolution imaging [44, 45], *in vitro* reconstitution assay [46, 47] and near-infrared deep tissue imaging [48, 49], and adoptions of the new developments of live-cell nano thermometry, membrane potential sensors and pH probes will together form a powerful platform for multifunctional imaging, sensing [50], therapy [48] and even tracking the pace of life [51] in living cells and organisms.

## Data Availability

The data that support the findings of this study are available from the corresponding author on reasonable request.

## Author Contributions

X.D., Q.P.S. and D.J. designed the project; X.D. prepared UCNPs and conjugates; M.M. synthesized the polymer; X.D., D.W., Y.L. and Q.P.S. built optical system, performed experiments and analyzed data; X.D., Q.P.S. and D.J. wrote the paper; Q.P.S. and D.J. supervised the studies.

## Acknowledgements

The authors acknowledge the financial support from the Australia National Health and Medical Council (NHMRC, APP1177374 to Q.P.S.), the Australia National Heart Foundation (NHF, 102592 to Q.P.S), the National Natural Science Foundation of China (NSFC, 61729501), the Major International (Regional) Joint Research Project of NSFC (51720105015), the Science and Technology Innovation Commission of Shenzhen (KQTD20170810110913065), the Australia China Science and Research Fund Joint Research Centre for Point-of-Care Testing (ACSRF658277, SQ2017YFGH001190), the Australian Research Council Laureate Fellowship Program (D.J., FL210100180) and the China Scholarship Council (no. 201706170028 to X.D. and no. 201706170027 to D.W.).

## Supporting Information

Materials and experimental procedures, bio-stability of nanoparticles, ATR-FTIR spectra, UV-Vis standard curve of TPP and data analysis process.

