## Supporting Information for "Spatiotemporally Mapping Thermodynamics of Lysosomes and Mitochondria using Cascade Organelle-Targeting Upconversion Nanoparticles"

for

**Materials and Methods Details**

**Reagents**

Yttrium(III) chloride hexahydrate (YCl_3_•6H_2_O, 99.99%), ytterbium chloride hexahydrate (YbCl_3_•6H_2_O, 99.99%), erbium chloride hexahydrate (ErCl_3_•6H_2_O, 99.9%), ethanol, cyclohexane, 1-octadecane (ODE), and oleic acid (OA) were purchased from Sigma-Aldrich. PEGMEMA_80_-*b*-EGMP_3_ di-block copolymer (polymer) was synthesized as previously described [1]. Poly-l-lysine (PLL), (3-carboxypropyl)triphenylphosphonium bromide (TPP), 1-Ethyl-3-(3-dimethylamino-propyl)-carbodiimide (EDC), N-Hydroxysuccinimide (NHS), MES buffer, HEPES buffer, Tetrahydrofuran (THF), N, N-Dimethylformamide (DMF), Bovine Serum Albumin (BSA), dimethyl sulfoxide (DMSO), carbonyl cyanide-4-(trifluoromethoxy)phenylhydrazone (FCCP) and ionomycin calcium salt were purchased from Sigma-Aldrich with reagent grade or higher. Dulbecco’s Modified Eagle Medium (DMEM), fetal bovine serum (FBS), Phosphate-Buffered Saline (PBS), Penicillin-Streptomycin (PS), LysoTracker (LysoTracker™ Deep Red FM, Invitrogen™ L12492) and MitoTracker (MitoTracker™ Deep Red FM, Invitrogen™ M22426) were purchased from Life Technologies (Thermo Fisher). Chloroquine (CQ, 14774S) was purchased from Cell Signaling Technology. Deionized (DI) water was generated by Milli-Q IQ Water Purification System (Millipore).

**Synthesis of hydrophilic upconversion nanoparticles (UCNPs)**

NaYF_4_: 20%Yb^3+^, 2%Er^3+^ nanoparticles were synthesized following the previously reported protocols with some modification [2, 3]. In a typical process, prepare stock solutions of YCl_3_ (0.4 M), YbCl_3_ (0.2 M) and ErCl_3_ (0.1 M) freshly. Then, 6 mL of OA and 15 mL of ODE were added into a 50 mL round bottom flask with three necks, and then added 1.95 mL of YCl_3_ stock solution, 1.0 mL of YbCl_3_ stock solution and 0.2 mL of ErCl_3_ stock solution. The mixture was stirred under argon protection and then heated to 100 °C for 10 minutes and 160 °C for 30 minutes to get rid of the methanol and H_2_O. Secondly, 5 mL of NaOH (0.1 g) - NH_4_F (0.14815 g) methanol solution was added into the mixture until it cooled down to 30 °C and stirred for 30 minutes. Subsequently, the mixture was heated to 100 °C for 30 minutes and then heated to 300 °C for 1.0 hours. Finally, 5 mL of ethanol was added into the mixture to precipitate UCNPs. UCNPs were washed 3 times with ethanol and cyclohexane before using. Finally, the UCNPs were dissolved in the 10 mL of cyclohexane and stored in the fridge.

To obtain hydrophilic UCNPs, the nanoparticles were first modified with PEGMEMA_80_-*b*-EGMP_3_ di-block copolymer (polymer) [4]. 5 mg of UCNPs and 5 mg polymer were dissolved in 1 mL of THF and then shaken at room temperature for 12 hours. Next, the UCNPs coated with polymers (UCNPs@polymer) were washed with THF three times and DI water three times. Finally, the UCNPs@polymer was dissolved in 0.5 mL of DI water and stored at 4 °C for further use.

**Bioconjugation of upconversion nanoparticles with (3-carboxypropyl)triphenylphosphonium bromide (TPP)**

To functionalize hydrophilic nanoparticles with amine groups, 0.5 mL of UCNPs@polymer were dissolved in the 0.5 mL of PLL solution and stirred at room temperature for 40 minutes. The reaction was stopped by centrifugation and washed three times with DI water. Finally, the UCNPs functionalized with amine groups (UCNPs@PLL) were stored at 4 °C for further use.

For the bioconjugation of UCNPs@PLL with TPP, 30 mg of TPP, 10 mg of EDC and 10 mg of NHS were first dissolved into 1mL of DMF by ultrasound. After 1 hour, 10 mg of UCNPs@PLL were added to the mixture. The reaction was stirred at room temperature for 12 hours and then washed with DMF and DI water. Finally, the product UCNPs@PLL conjugated with TPP (UCNPs@TPP) were stored in 0.5 mL of DI water at 4 °C for further use.

**Characterization**

The morphology of UCNPs was characterized using the FEI Tecnai transmission electron microscopy (TEM, FEI, U.S.A.). The hydrodynamic size and zeta potential of UCNPs were determined by a zeta sizer nano (Malvern, U.K.). The spectra were measured by a custom-built spectrometer.

**Cell Culture, Labelling and Colocalization of UCNPs with MitoTracker and/or LysoTracker**

The HeLa cells were seeded on the fluoro-dish (35 mm) at 10^5^ cell counting/density and incubated in the DMEM medium containing 10% v/v FBS for 12 hours. Then the cells were washed 3 times with PBS and incubated with UCNPs (50 µg/mL, 1 mL) for 12 hours at 37 °C with 5% CO_2_. Next, the cells were washed with PBS and incubated with 200 nM MitoTracker and/or 200 nM LysoTracker for 0.5 hours. Finally, the cells were washed with PBS and waiting for imaging in DMEM medium.

**Relative temperature sensing sensitivity and accuracy**

The relative temperature sensing sensitivity (S_R_) and accuracy (σT) are defined as follows [5]:

$$S_{R}= \frac{dRatio}{dT}\frac{1}{Ratio} (1)$$

$$\sigma T= \frac{\sigma R}{S_{R} \cdot R} (2)$$

where dRatio/dT can be obtained from the slope of the calibration curve represented in Figure 3B and σR is the standard deviation of the ratio. The ratio was obtained from the following equation (**Figure 3B**):

$$Ratio=10.3715T - 3.88731 (3)$$

**Temperature mapping and temperature sensing of mitochondria and lysosomes.**

The HeLa cells were cultured in DMEM containing 10% v/v FBS and 1% v/v Penicillin-Streptomycin at 37 °C with 5% CO_2_. Cells were transferred into the fluoro-dish (35 mm in diameter with No.1 coverglass bottom). Then the cells were incubated in the dish with 1 mL culture medium containing 50 µg/mL UCNPs@TPP or/and UCNPs@PLL at 37 °C for 12 hours. The UCNPs labeled cells were imaged on a custom-built total internal reflected fluorescent TIRF microscope with an external temperature controller by a 980 nm laser excitation under different temperatures (from 31 °C to 40 °C).

For the Chloroquine (CQ), carbonyl cyanide-4-(trifluoromethoxy)phenylhydrazone (FCCP), and Ca^2+^-induced temperature changes, first, the UCNPs@TPP were incubated with HeLa cells for 12 hours. Then, the HeLa cells were stained with MitoTracker and LysoTracker. Next, CQ (200 nM), FCCP (10 µM) or ionomycin calcium salt (1 µM) was incubated with HeLa cells. The fluorescence intensity ratio (I_525_ / I_545_) of UCNPs@TPP was determined with a 980 nm laser.

**Statistical Analysis.** The student’s t-test was applied to examine the differences among variables. Data were shown as mean ± SD. *p values ≤ 0.05 are considered to be statistically significant.

**Figure S1**

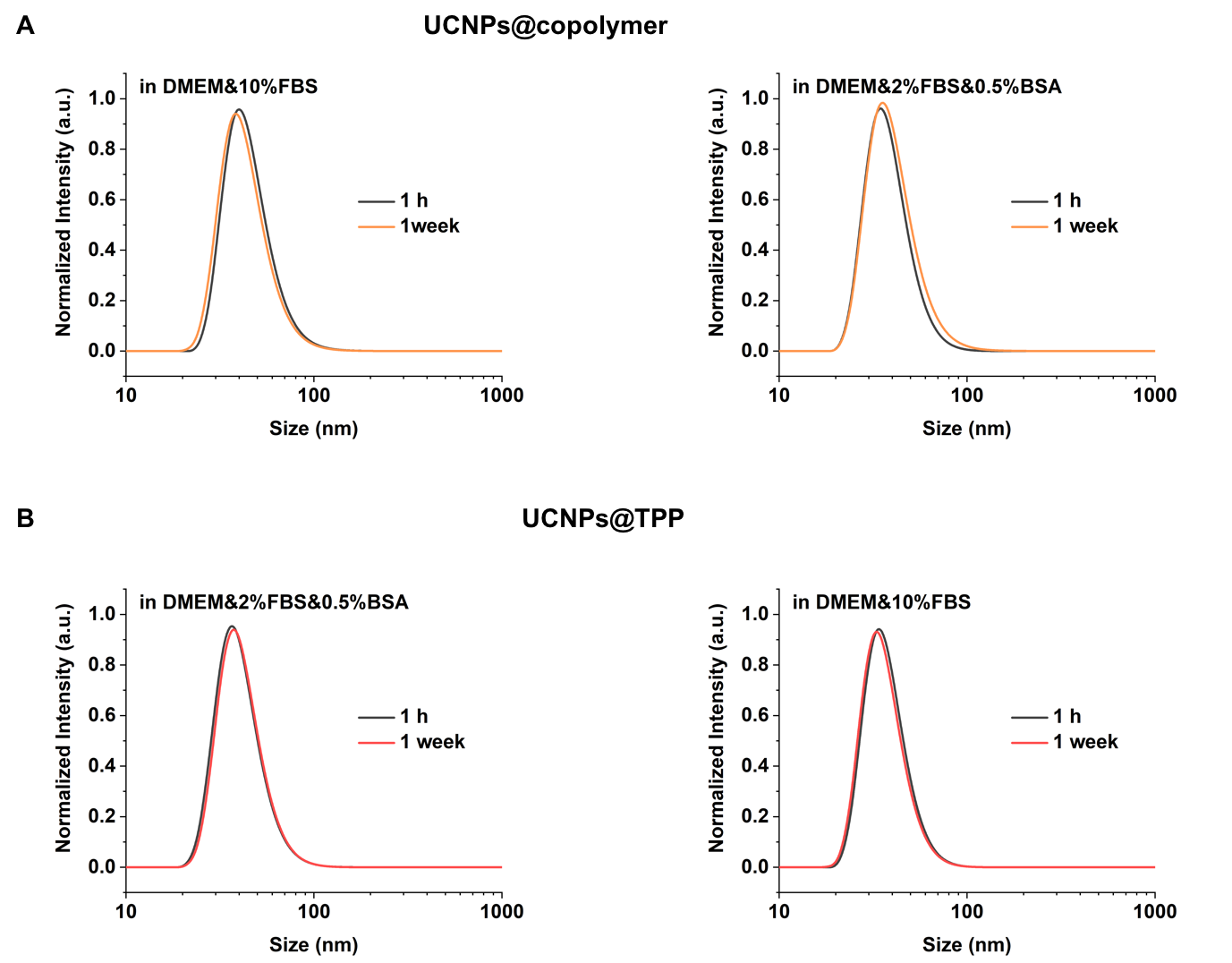

**Figure S1** The long-term bio-stability of UCNPs in aqueous solution. A. DLS results of UCNPs@copolymer in culture medium after 1 week. B. DLS results of UCNPs@TPP in culture medium after 1 week.

**Figure S2**

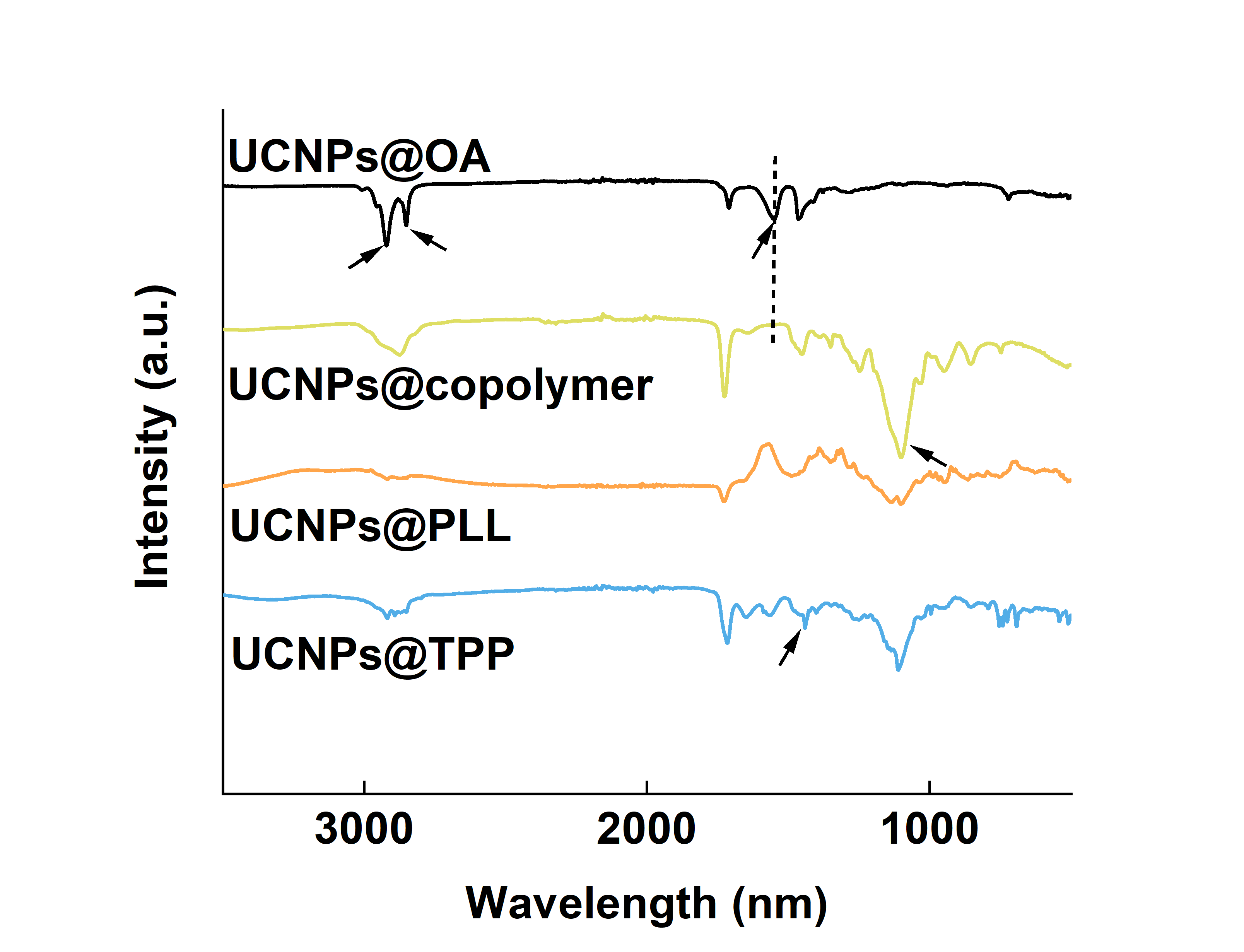

**Figure S2** The ATR-FTIR spectra of prepared UCNPs@OA, UCNPs@copolymer, UCNPs@PLL and UCNPs@TPP.

**Figure S3**

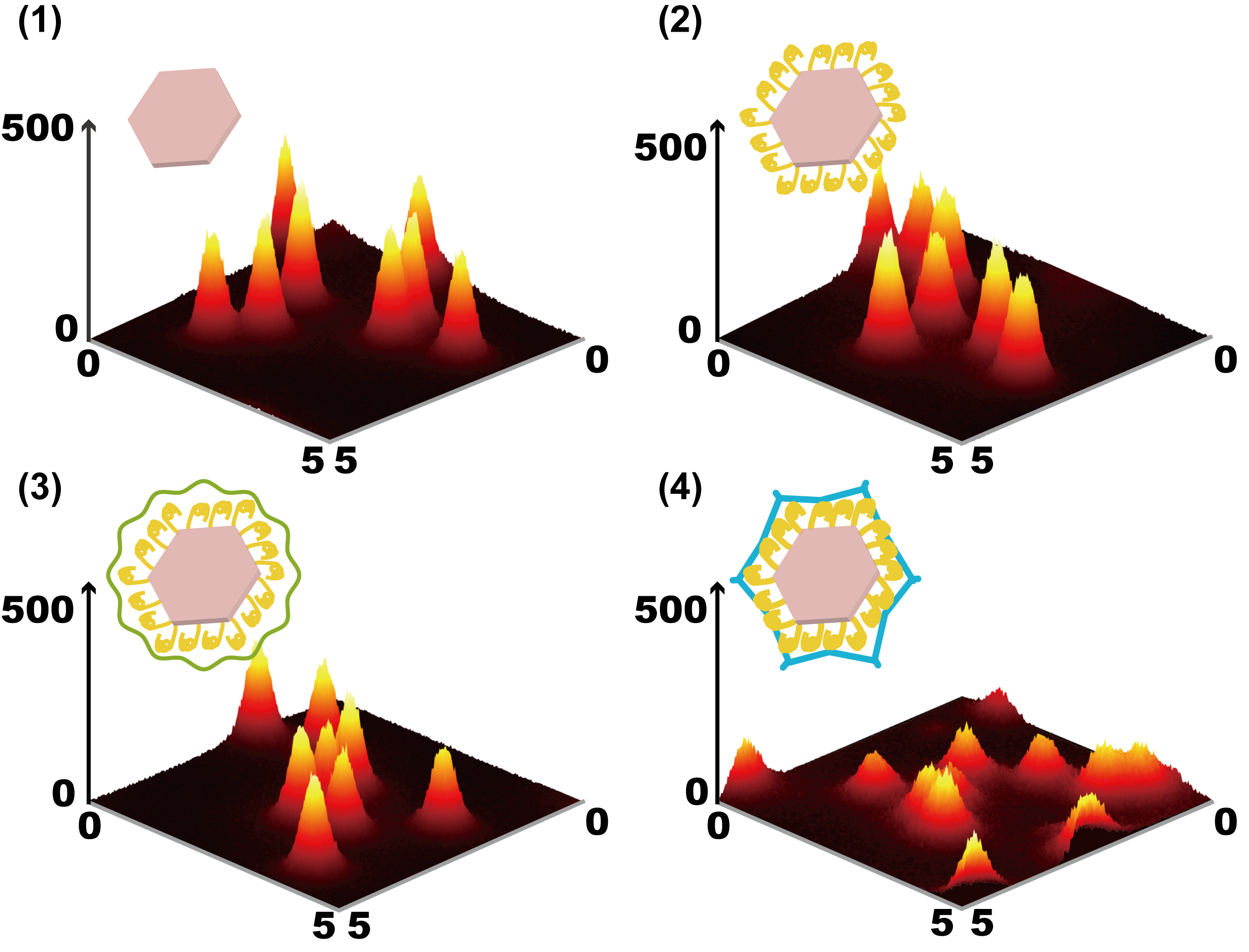

**Figure S3** Luminescence emission images from single UCNPs coated with OA, copolymer, PLL, and PEG under 980 nm excitation.

**Figure S4**

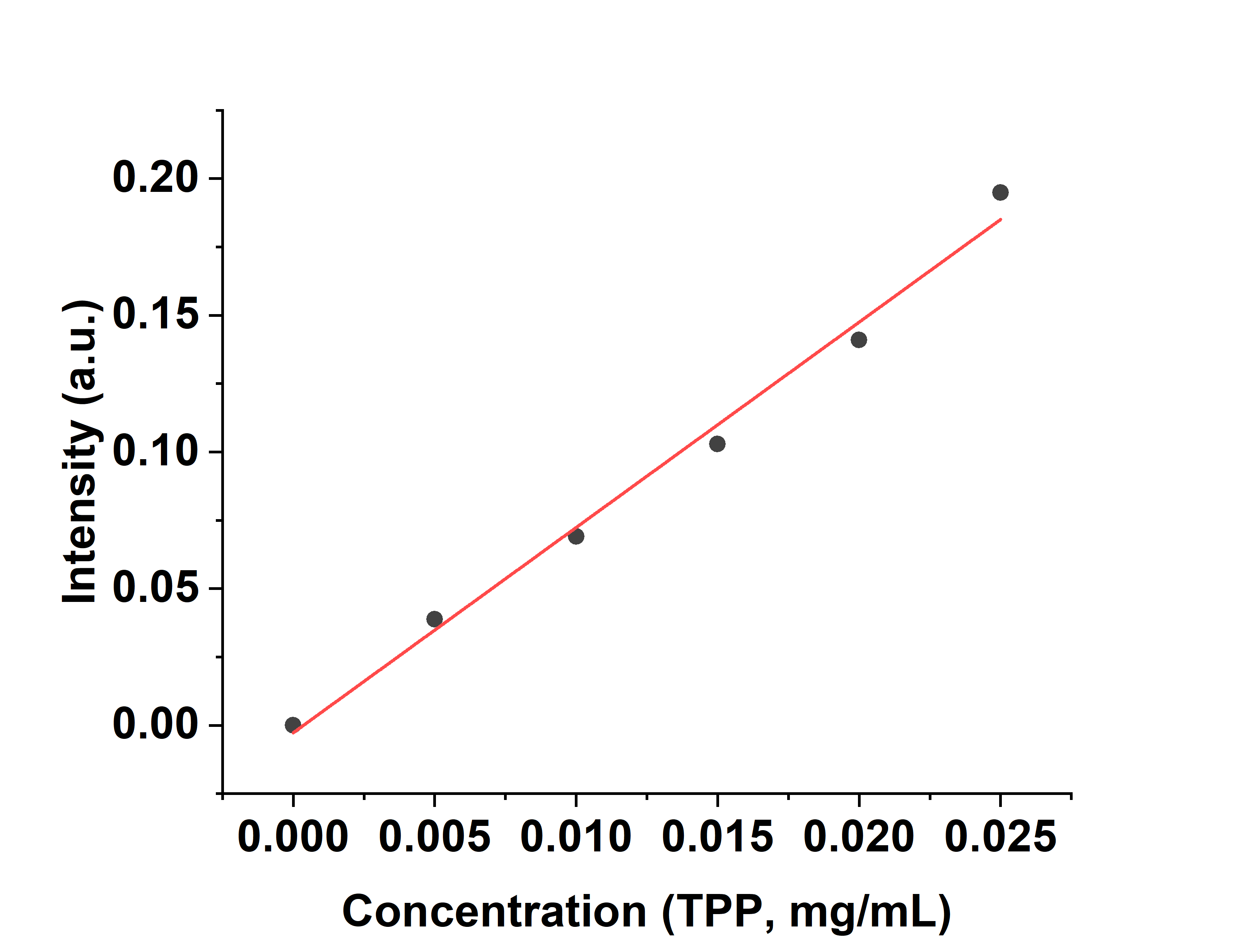

**Figure S4** UV-Vis absorption standard curve of TPP (268 nm) at various concentrations. It should be noted that for the TPP standard curve, as the EDC and NHS mixture in the sample solution showed obvious UV-Vis absorption, corresponding EDC and NHS need to be added as background into the standard TPP solution.
